# *In-vivo* evaluation of the anti-diarrheal effect of *Lactococcus lactis subspecies lactis* and *Lactococcus piscium* isolated from yogurt

**DOI:** 10.1101/2020.07.30.226688

**Authors:** Abu Sayeed Mohammad Mahmud, Mobarak Chowdhury, Rasheda Akter, Saiful Islam, Santosh Mazumdar, Tarannum Taznin, Rocky Chowdhury, Hridika Talukder, Habibur Rahman Bhuiyan

## Abstract

Lactobacillus and Lactococcus species found in the yogurt of different sources are most widely assayed and used all over the world as a probiotic agent. This study aimed to isolate and identify novel probiotic agents with therapeutic value against diarrhea. Initially, the probiotic properties of the isolated lactic acid bacteria from the yogurt samples of the Chittagong division, Bangladesh, were evaluated. All probiotic candidates inhibited the growth of selected pathogens, including *Escheriachia coli*, Serratia sp. *Salmonella paratyphi*, Streptococcus Group-B, *Staphylococcus aureus, Haemophillus influenzae, Bacillus subtillis*, and *Klebsiella pneumoniae. Lactococcus lactis* subsp lactis were found most useful in inhibiting all the selected pathogens. When the probiotics were applied against castor oil-induced diarrhea in the rat model, *Lactococcus lactis* subspecies *lactis* and *Lactococcus piscium* were found significantly effective relative to the controls indicating their potentiality as an alternative therapeutic against diarrhea.

**Highlights:**

1. *Lactococcus lactis* subspecies lactis and *Lactococcus piscium* has shown potentiality to be a therapeutic agent against castor-oil induced diarrhea in an animal model.
2. *Lactococcus lactis* subspecies lactis and *Lactococcus piscium* inhibited the growth of specified pathogens.

## Introduction

Probiotic bacteria obtained with food are frequently claimed to improve human health. From the view of the clinical field, probiotics are very beneficial for their unique therapeutic properties and inducing responsive actions. *In vitro* study revealed that the gut Lactic acid bacteria influence immunomodulation, induce the production of anti-inflammatory cytokines, and thus prevent the carcinogenesis [1]. A similar type of study revealed the ability of lactic acid bacteria (LAB) for removal of cholesterol, down-regulation of NPC1L1 protein, and deconjugation of bile salt [2]. Probiotics administration also significantly reduce bone loss (p < .0001) and gingival inflammation (p < .0001) [3].

In the rabbit model, *Lactococcus lactis* protect the rabbit against the development of necrotizing enterocolitis in the presence of *Cronobacter sakazakii* [4]. In animals, Lactobacillus sp. alone or with the combination of other probiotics play a role in the eradication of *Helicobacter pylori* growth and inflammatory infection [5]. Probiotics have the potentiality to modify the natural history of inflammatory bowel disease (IBD) according to the data derived from animal models [6]. In the animal model, the administration of probiotics improves clinical symptoms, mucus production, and histological alterations [7]. In rats, Bifidobacterium and Lactobacillus have been shown to influence phagocytosis of macrophages, cytotoxicity of natural killer cells, and mediate adaptive immunity [8]. *L. plantarum* MG989 and *L. fermentum* MG901 have shown the potentiality to inhibit the yeast growth, and thus helping to clear vulvovaginal candidiasis *in vivo* [9]. Besides, pro-and prebiotics has been found to revive healthy digestive system function in autistic patients [10].

In society, infectious diseases are the most critical problem with significant morbidity, and mortality occurs worldwide due to gastrointestinal infections [11]. It is reported in a rural Thai population that probiotic Bacillus bacteria abolish the pathogen *Staphylococcus aureus* [12]. In the gut, Bacillus sp has been shown to produce biofilm biomass and exert antimicrobial and enzymatic activity, thus protect from GIT and other infections [13]. Probiotics are typically involved in the production of bacteriocin and bacteriocin like substances that inhibit the growth of various spoilage and pathogenic microorganisms, including Streptococcus Group-B, *Bacillus aureus, Escherichia coli, Salmonella paratyphi, Pseudomonas aeruginosa, Serratia* sp., *Klebsiella pneumonia*, and *Neisseria meningitides* [14]. *In-vitro* antimicrobial activities of a *Lactobacillus sp.* against diverse enteric bacteria, including *Vibrio cholerae*, was demonstrated [14]. A probiotic like Bacillus strains are found to inhibit the growth of *E. cecorum in vitro* [15]. Probiotic *L. plantarum* ATCC 8014 has shown the potential for controlling the growth and infections by Clostridium sp [16]. It has demonstrated *in vitro* experiment that *Streptococcus dentisani* attached to gingival cells can inhibit periodontal pathogens [17].

Though one of the criteria of being a probiotic is not carrying transmissible antibiotic resistance genes [18], the characterization of the members of the genera *Lactococcus* and *Lactobacillus*, particularly in terms of antimicrobial resistance, is often neglected as they have a long history of being the most commonly given generally-recognized-as-safe (GRAS) status. In contrast, members of the other genera of LAB contain some opportunistic pathogens. In many cases, antibiotic resistances of LAB are intrinsic, which are not transmissible. Besides, they are sensitive to many clinically used antibiotics, even in the case of a LAB-associated opportunistic infection. Thus, there is no safety concern with intrinsic resistance [19]. Preferably, inherent resistance of LAB against some antibiotics can be utilized for selective combinatorial therapy of antibiotics and probiotics. In a study, 55 probiotic strains of different bacteria were isolated from commercial dairy and pharmaceutical products. Where, all the isolated strains showed resistance to aztreonam, cycloserine, kanamycin, nalidixic acid, polymyxin B, and spectinomycin. On the other hand, they showed susceptibility to ampicillin, clindamycin, bacitracin, dicloxacillin, novobiocin, erythromycin, penicillin G, and rifampicin [18]. Similarly, a different group investigated antibiotic resistance of 41 strains of LAB where most of them were resistant to amikacin, ciprofloxacin, gentamycin, and trimethoprim/sulfamethoxazole; but were sensitive against chloramphenicol, ampicillin, tetracycline, amoxicillin, imipenem, and cephalothin [20].

Probiotics are widely present in a variety of foods including dairy-based products such as acidophilus milk, acid-whey, ice-cream, lassi, cheese, curd, non-fermented goat’s milk beverage, frozen symbiotic dessert, and yogurt as well as present in non-dairy sources including products from cereals, fruits, vegetables, meat, fishes, and soy [21]. Though there are both non-dairy and dairy sources of probiotics, many studies have shown that fermented dairy products are the best matrices to deliver probiotics [22]. Previously, 21 different probiotic lactic acid bacteria were isolated from yogurt of the local market of the Chattogram Division, Bangladesh [14]. In Spain, a group of researchers demonstrated the prevalence of LAB in commercial dairy products ranging from 10^8^ – 10^9^ CFU g^−1^ [23]. In a separate study conducted in Mali found the incidence of LAB in local fermented-dairy product around 10^8^ CFU mL^−1^ in all-season [24]. Also, probiotic properties-bearing LAB was isolated from fermented dairy milk and fermented olives in Thailand and Greece, respectively [25, 26].

In Bangladesh, yogurt is one of the oldest fermented milk products known and consumed by large sectors of the population as a part of their daily diet. In most of the areas of Bangladesh, different types of traditional yogurts are found, but their probiotic role was not studied. Previously, the probiotic properties of *Lactobacillus spp.* and *Lactococcus spp.* were examined by *in vitro* experiments [14, 27]. However, the probiotic activities of lactic acid bacteria did not observe for Bangladeshi isolates collected from yogurt. Therefore, this study aimed to evaluate the utilization of anti-diarrheal probiotic for cholera patients to determine if this intervention can successfully decrease watery diarrhea in the treated mouse.

## Materials and method

### Determination of optimum pH and temperature for efficient antimicrobial activity

The De Man, Rogosa, and Sharpe (MRS) broth (Oxoid) was adjusted at different pH (pH 3, 3.5, 4, and 4.5) using 0.1 N Acetic acid and 0.1 N NaOH. Aliquots of 10 ml medium from each pH set were dispensed in a separate test tube and autoclaved. Four sets of tubes were inoculated with culture suspension of five isolates and incubated at 27 °C, 37 °C and 45 °C temperature for 24-48 hours. Using a spectrophotometer at 560 nm, the turbidity was measured and filtered with the help of the Whatman filter paper (Whatman International Ltd. England). Measuring pH, the culture filtrate (Hanna Instrument Ltd. U.K.) was applied for antimicrobial activity against the respective pathogenic bacteria by the agar well diffusion method.

### Bile salt tolerance of the isolates

With different concentrations (i.e., 1.0%, 2.0%, and 3.0%) of bile salt, MRS broth was prepared and dispensed at 10 ml per tube with the selected isolates. The incubation condition was 37 ± 1 °C for 24-48 hours. Using the pour plate method, 100 μl of culture from each concentration was grown in agar medium at 37 °C for 24 hours for comparative growth.

### Activities of selected lactic acid against specified pathogens

The procedure for observing the activity of selected lactic acid bacteria against pathogens developed previously. Briefly, an overnight grown pathogenic bacteria culture was suspended in 2 ml of sterilized saline water and mixed thoroughly. A total of seven pathogenic microorganisms were used in this method, namely *Bacillus subtilis, Staphylococcus aureus. Streptococcus Group-B, Escherichia coli, Serratia sp., Klebsiella pneumonia*, and *Haemophillus influenzae*. Growth inhibition was done in Mueller Hinton by the Well diffusion method, and media were seeded uniformly with 2.7×10^3^ cells per ml of test organisms and incubated at 37 °C for 24 hours. The anti-bacterial activity of the test agent was determined by measuring the zone of inhibition expressed in mm in diameter.

### Assay of Antibiotic Sensitivity Pattern

To determine the antibiotic sensitivity pattern, the disk diffusion method was followed. In this method, Mueller Hinton plates were prepared and swabbed with the suspension of selected isolates using a sterile cotton bud. The antibiotic disks were placed on the surface of the plate, maintaining equal distance and placed at 4°C for 1-2 hours for proper diffusion of antibiotics followed by incubation for 18-24 hours at 37°C. The zone of inhibition measured by the zone of diameter for antibiotic sensitivity or resistance [28].

### *In-vitro* testing for the anti-diarrheal effect of selected isolates

The *in vivo* anti-diarrhoeal activity of the Castor oil-induced diarrheal model was performed by [29] suggested method. Thirty-five Wistar rats were randomly divided into seven equal groups (n=5), Control group, positive control group, and five treated groups. Control group were received only distilled water 2 ml/rat, the positive control group received Loperamide (Opsonin, Bangladesh) 4.17 mg/kg as standard and treated groups received selected isolates or probiotics at the dose of 10^6^CFU/ML. Rats were encased in separate cages having paper placed below for the collection of fecal matters. Diarrhea was induced in rats by oral administration of castor oil (2.0 ml/rat). The anti-diarrheal activity was determined by giving probiotics and drugs orally 1 hour before the administration of a standard dose of 2.0 ml of castor oil. The number of both hard and soft pellets was enumerated at every hour over 6 hours period for each rat. Diarrhea was confirmed as the presence of stool with a fluid material that stained the paper placed beneath the cages.

### Statistical analysis

All the statistical values of anti-diarrheal, tests were reported as mean + SEM (Standard error of the mean). Statistical differences between the mean of the various groups were analyzed using the Student’s “t” test. Probability (p) value of 0.05 or 0.01 was considered significant. All the graphical presentations and statistical calculations were prepared using “Microsoft Excel-2007”. Mean values were considered significantly different if *P*< 0.05, 0.01.

## Results

### Bile salt tolerance

As probiotic organisms must tolerate varying concentrations of bile salt in the intestinal tract, the isolates were tested for their sensitivity to different concentrations of bile salt (i.e., 1%, 2%, and 3%). All the selected strains showed the highest number of the colony (cfu/0.1ml) for 1% bile salt concentration and lowest number of the colony for 3% bile salt concentration, which was 120 and 85 for *Lactococcus lactis* subsp. *Lactis*, 110 and 45 for *Lactococcus raffinolactis*, 80 and 40 for *Lactococcus piscium*, 360 and 180 for *Lactobacillus delbrueckii*, and 510 and 350 for *Lactobacillus plantarum*, respectively. With the increase of bile salt concentration, the number of colonies reduced for all the selected isolates, which tolerate up to 3% bile salt. The ability of isolates to tolerate varying concentrations of bile salt showed in **Table 1**.

**Table 1:** Bile salt tolerance of the selected isolates

| Isolates | No. of colony grown on the plates (cfu/100 µL) |  |  |
| --- | --- | --- | --- |
|  | 1.0% | 2.0% | 3.0% |
| *Lactococcus lactis* subsp. *lactis* (LcL) | 120 | 100 | 85 |
| *Lactococcus raffinolactis* (LcR) | 110 | 60 | 45 |
| *Lactococcus piscium* (LcP) | 80 | 55 | 40 |
| *Lactobacillus delbrueckii* (LbD) | 360 | 220 | 180 |
| *Lactobacillus plantarum* (LbP) | 510 | 450 | 350 |

### Assay of Antibiotic Susceptibility pattern

The antibiotic sensitivity pattern of the selected isolates was also determined to investigate the severe effect of any antibiotics against them. The application of this type of antibiotic with probiotic organisms may demolish the therapeutic efficacy of the probiotic. The Antibiogram obtained is shown in figure 2. It can be demonstrated from the graph that isolates LcL showed 50% resistance, 37.5% sensitivity, and 12.5% intermediate among 16 antibiotic disks. LcR had 25% resistance, 62.5% sensitive, and 12.5% intermediate among 16 antibiotic disks. Isolate LcP revealed 18.75% resistance, 56.25% sensitivity, and 25% intermediate among 16 antibiotic disks and 80% resistance, 20% sensitivity, and no intermediate found among 21 antibiotic disks for LbD. The LbP isolate showed 20% resistance, 71% sensitivity, and 9% intermediate among 21 antibiotic disks.

**Fig. 1.**
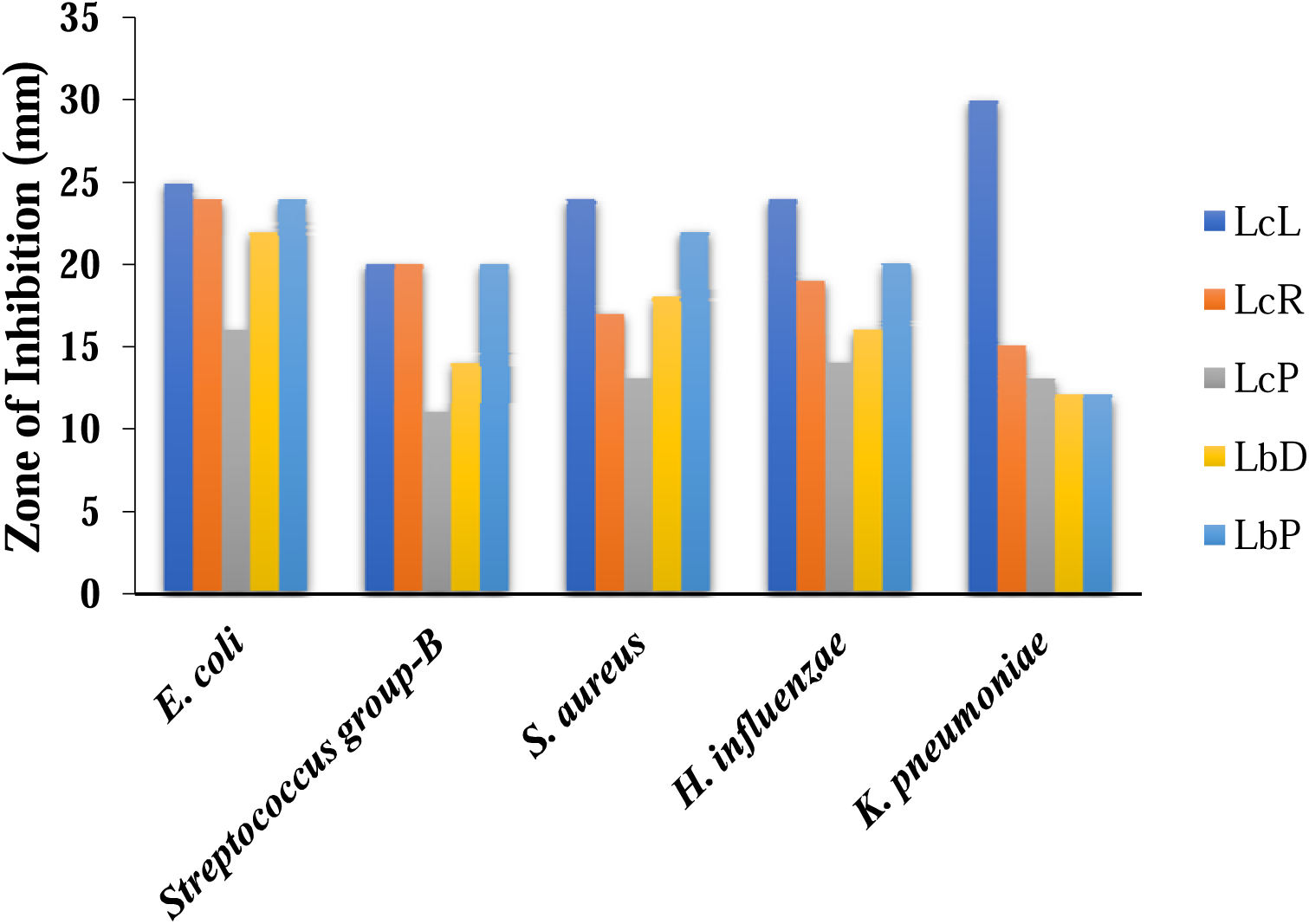
Zone of inhibition produced due to the activity of bacteriocin produced by isolate LcL, LcR, LcP, LbD, and LbP against five pathogenic organisms. *Isolates (LcL: *Lactococcus lactis* subsp. *lactis*; LcR: *Lactococcus raffinolactis;* LcP: *Lactococcus piscium;* LbD: *Lactobacillus delbrueckii;* LbP: *Lactobacillus plantarum*

**Fig. 2:**
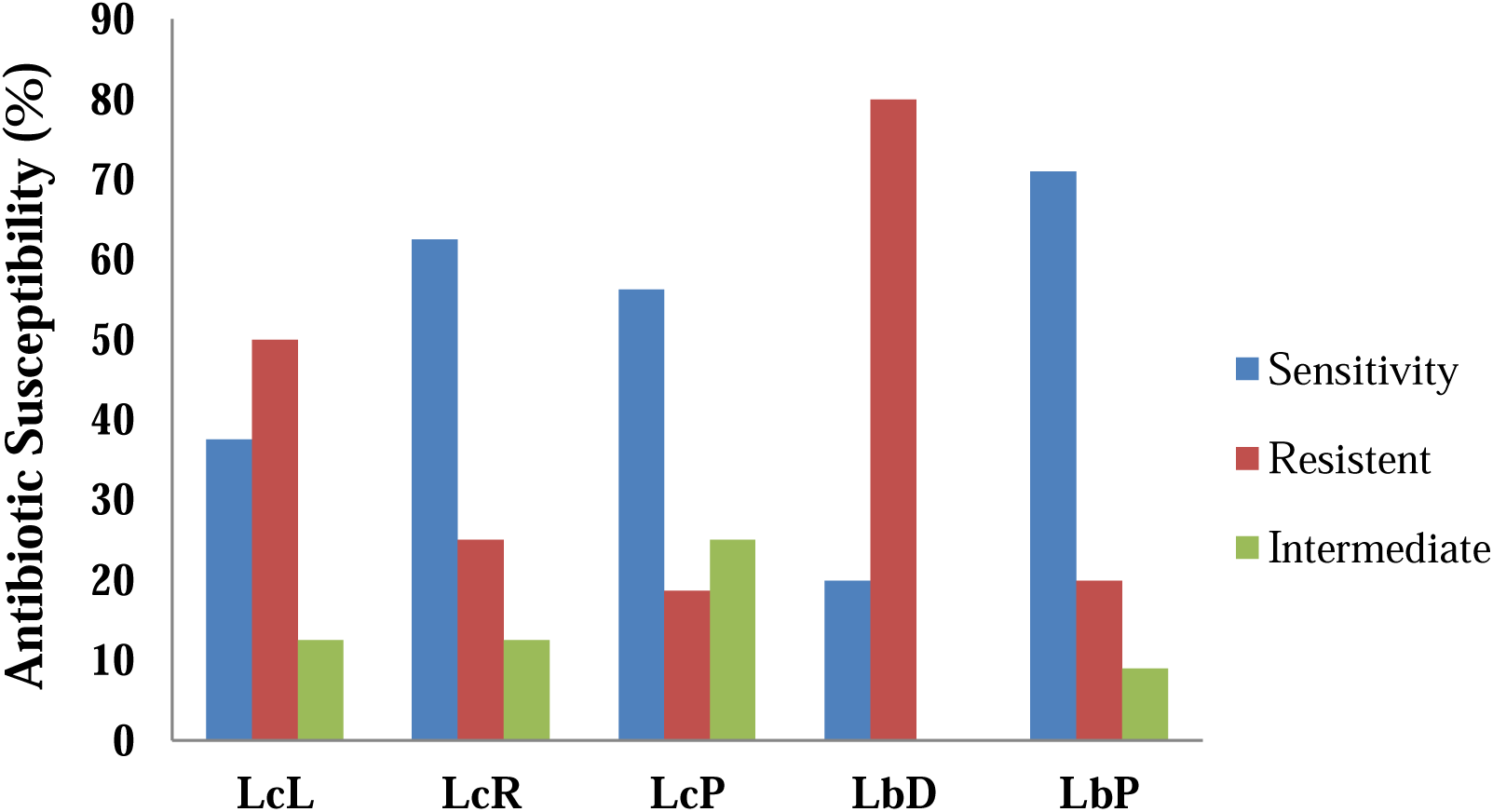
Percentage of antibiotic resistance, intermediate and sensitivity among the five Isolates. A total of 16 antibiotic disks were applied against *Lactococci sp*. and 21 for *Lactobacilli sp*.

### Anti-bacterial activities of isolates

Lactic acid bacteria produce some antimicrobial substances. One of the substances is bacteriocin, which has emerged as an essential antimicrobial substance. The antimicrobial activity of bacteriocin was examined against the microorganisms’ *Escherichia coli, Streptococcus Group-B, Staphylococcus aureus, Haemophillus influenza, Klebsiella pneumonia*. LcR, LcP, LbD, and LbP produced a zone of inhibition (mm) respectively 25, 24, 16, 22 and 24 mm against pathogenic bacteria *E. coli*, and 20, 20, 11, 14, 20 mm against pathogenic bacteria *Streptococcus Group-B*, and 24, 17, 13, 18, 22 mm against pathogenic bacteria *Staphylococcus aureus*, and 24, 19, 14, 16, 20 mm against pathogenic bacteria *Haemophillus influenza*, and 30, 15, 13, 13, 12 mm against pathogenic bacteria *Klebsiella pneumonia.* The measured diameter of inhibition zones accomplished that all the isolates have anti-bacterial effects against the pathogenic microorganisms **(Figure 1).**

### Effect of *Lactococcus* species against Castor Oil induced diarrhea

The *Lactococcus* species was evaluated in castor oil-induced diarrheal rats as compared to normal rats. The feces at sixth hours for *Lactococcus* species treated group at 10^6^CFU/ML were significantly (p<0.05, p<0.01) decreased as compared to the control group **(Table 4)**.

**Table 2:**
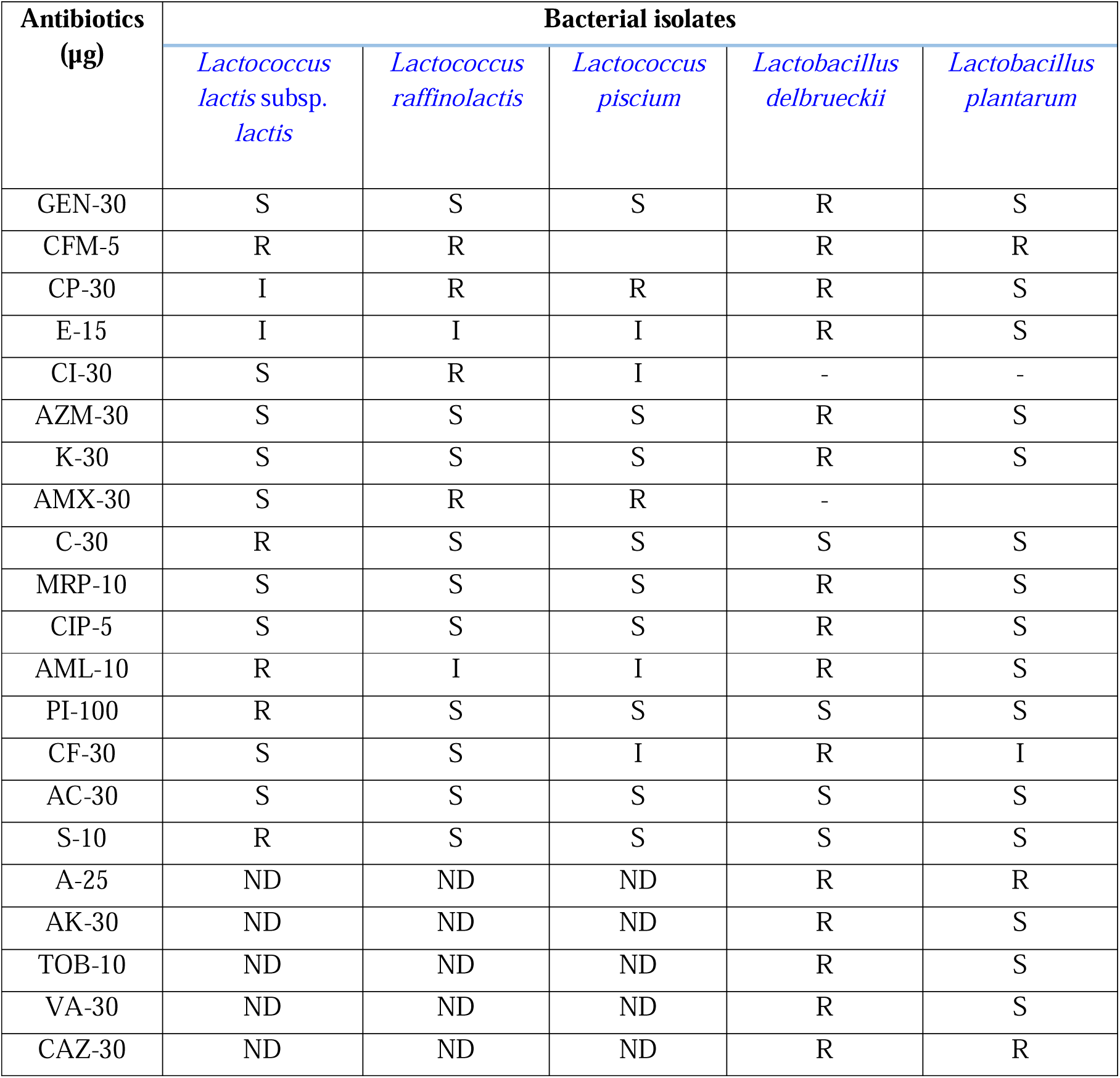
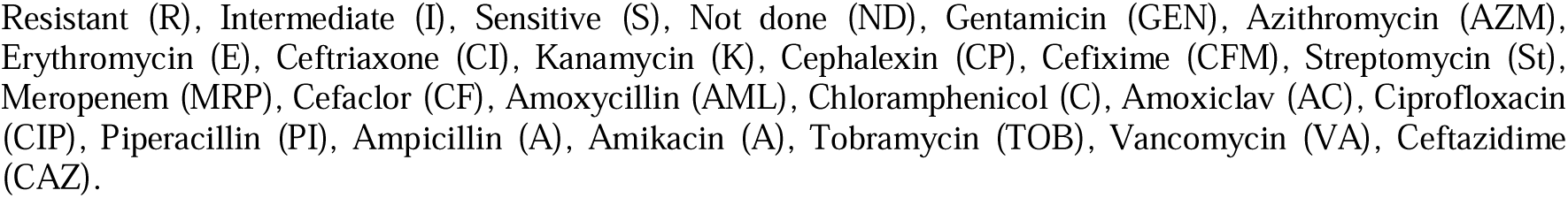
Antibiotic sensitivity pattern of the selected isolates

| Antibiotics<br>(µg) | Bacterial isolates |  |  |  |  |
| --- | --- | --- | --- | --- | --- |
|  | *Lactococcus lactis* subsp. *lactis* | *Lactococcus raffinolactis* | *Lactococcus piscium* | *Lactobacillus delbrueckii* | *Lactobacillus plantarum* |
| GEN-30 | S | S | S | R | S |
| CFM-5 | R | R |  | R | R |
| CP-30 | I | R | R | R | S |
| E-15 | I | I | I | R | S |
| CI-30 | S | R | I | - | - |
| AZM-30 | S | S | S | R | S |
| K-30 | S | S | S | R | S |
| AMX-30 | S | R | R | - |  |
| C-30 | R | S | S | S | S |
| MRP-10 | S | S | S | R | S |
| CIP-5 | S | S | S | R | S |
| AML-10 | R | I | I | R | S |
| PI-100 | R | S | S | S | S |
| CF-30 | S | S | I | R | I |
| AC-30 | S | S | S | S | S |
| S-10 | R | S | S | S | S |
| A-25 | ND | ND | ND | R | R |
| AK-30 | ND | ND | ND | R | S |
| TOB-10 | ND | ND | ND | R | S |
| VA-30 | ND | ND | ND | R | S |
| CAZ-30 | ND | ND | ND | R | R |

Resistant (R), Intermediate (I), Sensitive (S), Not done (ND), Gentamicin (GEN), Azithromycin (AZM), Erythromycin (E), Ceftriaxone (CI), Kanamycin (K), Cephalexin (CP), Cefixime (CFM), Streptomycin (St), Meropenem (MRP), Cefaclor (CF), Amoxycillin (AML), Chloramphenicol (C), Amoxiclav (AC), Ciprofloxacin (CIP), Piperacillin (PI), Ampicillin (A), Amikacin (A), Tobramycin (TOB), Vancomycin (VA), Ceftazidime (CAZ).

**Table 3:** Assay of bacteriocin activity produced by the selected isolates

| Isolates | Pathogenic bacteria | Zone of inhibition (mm) |
| --- | --- | --- |
| ***Lactococcus lactis* subsp. *lactis* (LcL)** | *Escheriachia coli* | 25 |
|  | *Serratia sp.* | 20 |
|  | *Salmonella paratyphi* | 22 |
|  | *Streptococcus Group-B* | 20 |
|  | *Staphylococcus aureus* | 24 |
|  | *Haemophilus influenzae* | 24 |
|  | *Bacillus subtilis* | 18 |
|  | *Klebsiella Pneumoniae* | 30 |
| ***Lactococcus raffinolactis* (LcR)** | *Escheriachia coli* | 24 |
|  | *Serratia spp.* | 17 |
|  | *Streptococcus Group-B* | 20 |
|  | *Staphylococcus aureus* | 17 |
|  | *Haemophilus influenzae* | 19 |
|  | *Bacillus subtilis* | 18 |
|  | *Klebsiella Pneumoniae* | 15 |
| ***Lactococcus piscium* (LcP)** | *Escheriachia coli* | 16 |
|  | *Serratia sp.* | 20 |
|  | *Salmonella paratyphi* | 20 |
|  | *Streptococcus Group-B* | 11 |
|  | *Staphylococcus aureus* | 13 |
|  | *Haemophilus influenzae* | 14 |
|  | *Klebsiella Pneumoniae* | 13 |
| ***Lactobacillus delbrueckii* (LbD)** | *Escheriachia coli* | 22 |
|  | *Streptococcus Group-B* | 14 |
|  | *Staphylococcus aureus* | 18 |
|  | *Haemophilus influenzae* | 16 |
|  | *Klebsiella Pneumoniae* | 12 |
| ***Lactobacillus plantarum* (LbP)** | *Escheriachia coli* | 24 |
|  | *Streptococcus Group-B* | 20 |
|  | *Staphylococcus aureus* | 22 |
|  | *Haemophilus influenzae* | 20 |
|  | *Klebsiella Pneumoniae* | 12 |

**Table 4:** Effect of *lactic acid bacteria* species against castor oil-induced diarrhea.

| Groups | Treatment | Dose | Total Number of Faeces per hours | Total Number of Wet faces per hours |
| --- | --- | --- | --- | --- |
| Control | Distilled Water | 2ml/Rat | 33.25±3.28 | 28.75±2.56 |
| Positive Control | Loperamide | 4.17mg/kg | 9.75±0.48** | 3.25±0.48** |
| Treated | *L. lactis* subsp. *lactis* | 10<sup>6</sup>CFU/ML | 8.75±0.48** | 4.75±0.75** |
|  | *L. piscium* |  | 11.5±0.65** | 6.5±1.04** |
|  | *L. raffinolactis* |  | 20.5±1.55* | 11±1.08* |
|  | *L. plantarum* |  | 16.75±0.48* | 8.25±0.48** |
|  | *L. delbrueckii* |  | 26.25±1.38<sup>ns</sup> | 16.25±1.03* |

Values are expressed as mean ± SEM (Standard Error of the Mean). Statistical differences between the mean of the various groups were analyzed by using Student's t-test. \*: P <0.05; \*\*: P <0.01 mean significant difference compared positive control and treated with control. ns means not-significant. All the graphical presentation and statistical calculations were prepared using 'Microsoft Excel-2007'.

## Discussion

Acute diarrhea is one of the significant reasons for childhood morbidity as well as a load on society, which comes with an economic and emotional burden for the families of patients. Different kinds of medications, including those affecting ion transport as well as intestinal motility, adsorptive moieties, and living bacteria, have been applied to remediate acute diarrhea [30]. Our study assessed the potentiality of some selected LAB against castor oil-induced diarrhea in the mouse model.

The bile salt-hydrolyzing ability of probiotics has often been one of the criteria to select probiotic strain. Besides, a different kind of bile salt hydrolases are responsible for the catalysis of conjugated bile salt, have been found and characterized [31]. Previously, we have isolated five different lactic acid bacterial species bearing probiotic properties from yogurt samples of the Chittagong Division, Bangladesh [14]. In this study, the selected probiotics were grown in MRS broth containing varying concentrations of bile salt (1% – 3%) to check their ability to survive inside the gastrointestinal tract containing bile salt as bile salt has antimicrobial property [32]. Although all our probiotic isolates can tolerate bile salt concentration up to 3%, grow best at 1% **(Table 1)**. The results of this study are in conjunction with the previous research where it has been shown that *L. plantarum, L. casei, L. helvecticus* can tolerate bile salt 2% and best tolerated at 1% [33]. Likewise, in a different study, four strains of *Lactobacillus* were examined for their tolerance in different bile salt concentrations, where all of the isolates showed similar results [34]. The result indicates that our selected probiotics might be able to produce bile salt hydrolase (BSH), making them eligible for sustaining in the gut environment against the toxic effects of bile salt.

The probiotic LAB exerts its antimicrobial effects partly by secreting antimicrobial compounds, like bacteriocin, hydrogen peroxide, and organic acids. It has been proven that probiotics are effective against *Clostridium difficile*, which is responsible for *C. difficile*-associated diarrhea (CDAD) and proposed as a promising treatment option for *C. difficile* infection (CDI) [35]. Here, we have observed the antimicrobial activities of the probiotic isolates against 7 different specified pathogens. *Lactococcus lactis* subsp. *lactis, Lactococcus piscium, Lactococcus raffinolactis, L. delbrueckii*, and *L. plantarum* were found to inhibit the pathogenic bacteria, including *E. coli, Streptococcus Group-B, Staphylococcus aureus, Haemophillus influenza, Klebsiella pneumonia, Salmonella paratyphi*, and *Bacillus subtillis* **(Table 3)**. Consistent with our study, *L. plantarum* has shown higher antimicrobial activities against *E. coli* and *Staphylococcus aureus* than the others [36]. It has also been reported that *Lactococcus lactis sp. lactis* antagonistic against *Salmonella* sp. and *E. coli* strains, which support our findings [36]. Besides, like our finding, *L. piscium* found to have antimicrobial activities against various pathogens, including *E. coli* and *S. aureus* in other studies [37]. Similarly, *L. delbrueckii* showed significant anti-bacterial effects against *E.* coli and *S. aureus* [38, 39]. In addition, *L. plantarum* reported producing diverse types of bacteriocins that are effective against a vast number of bacterial species, including *E.* coli, *S. aureus*, and *Streptococcus* spp [40]. Our results indicate that the selected strains may be able to produce antimicrobial compound(s), making them competent for further experiments.

LAB may have intrinsic resistance against some antibiotics [18]. Intrinsic resistance of LAB to antibiotics could be beneficial to use the isolates *Lactococcus* & *Lactobacillus sp*ecies as probiotics with the combination of certain antibiotics. When selected, probiotics were exposed to 16 different antibiotics, *Lactococcus lactis* subsp. *lactis* was found sensitive to Gentamicin, Kanamycin, Azithromycin, Meropenem, Ciprofloxacin, Amoxyclav, and intermediate to Cephalexin, Erythromycin; but resistant to Ceftriaxone, Cefixime, Amoxicillin, Chloramphenicol, Piperacillin, Cefaclor, Streptomycin. Besides, *L. piscium* was found to be sensitive to Gentamicin, Azithromycin, Kanamycin, Chloramphenicol, Meropenem, Ciprofloxacin, Piperacillin, Amoxyclav, and Streptomycin, intermediate to Erythromycin, Ceftriaxone, Amoxicillin, and Cefaclor resistant to Cefixime, Cephalexin, amoxicillin. We found that *L. raffinolactis* is sensitive to Gentamicin, Azithromycin, Kanamycin, Chloramphenicol, Meropenem, Ciprofloxacin, Amoxyclav, and Streptomycin, intermediate to erythromycin, and resistant to Cephalexin and Amoxicillin **(Table 2)**. It has been reported that most *Lactococcus* species exhibit intrinsic resistance to trimethoprim, metronidazole, and cefoxitin, and the aminoglycosides kanamycin and gentamicin. Also, *L. lactis* found to carry antibiotic resistance genes erm(B), *tet*(M), tet(S), and *dfr*(A) [41]. In a different study, three multi-drug resistance genes, namely *lmr*(A), *lmr*(D), and *lmr*(P), were found in *L. lactis* [42]. Previously, *L. raffinolactis* found to be not resistant to Ciprofloxacin, Erythromycin, and Gentamicin, which matches our findings [43].

In our study, *Lactobacillus delbrueckii* showed sensitivity to Chloramphenicol, Piperacillin, Streptomycin, and Amoxiclav, resistance to all the other antibiotics we used for this isolate. At the same time, *L. plantarum* was resistant to Cefixime and Ampicillin, intermediate to Cefaclor, and sensitive to different antibiotics subjected to this strain **(Table 2)**. Similarly, various studies have shown that an increasing number of foodborne *Lactobacillus* species contain one or more antibiotic resistance genes [41]. Moreover, the presence of tet(M), *aph* (3’)-III, and *ant* (6) antibiotic resistance genes in *L. delbrueckii* and *erm*(B), erm(C), *tet*(M), *tet*(S), *tet*(W) in *L. plantarum* has been reported by researchers [44]. Previously, *L. delbrueckii* showed resistance against Vancomycin, Ciprofloxacin, Gentamicin, Tobramycin, and Nitrofurantoin. On the other hand, *L. plantarum* was not resistant against Meropenem, Ciprofloxacin, Gentamicin, Streptomycin, Chloramphenicol, Erythromycin, and Nitrofurantoin [45]. Our results support the previous findings.

Though all of our selected isolates were susceptible to some antibiotics resistance to other, three isolates of genus *Lactococcus* showed the lesser difference between the percentage of antibiotics that they were resistant and the number of antibiotics that they were susceptible **(Figure 2)**. Preferably, these isolates are better choices as they give us plenty of options for both combinatorial treatment and treatment for LAB-associated opportunistic infection, which usually does not occur for *Lactococcus* and *Lactobacillus* [19]. The resistance of the probiotic strains to some antibiotics could be used for both preventive and therapeutic purposes in controlling intestinal infections [46].

Finally, the selected *Lactococcus* strains were subjected to test their ability to ameliorate the diarrheal condition where we have induced rat with castor oil to produce diarrhea. We treated rats with both standard Loperamide and our isolated *Lactococcus* species and compared the number of feces and wet feces per hour between controls and treatments **(Table 4)**. In comparison to Loperamide, *Lactococcus* isolates, especially *Lactococcus lactis* and *Lactococcus piscium*, have the same effects on the rat. In a previous study, it was suggested that the LAB could exert their beneficial effects against diarrheal diseases by multiple mechanisms [47]. Probiotic bacteria, including *Lactococcus* spp. And *Lactobacillus* spp. showed a significant prevention effect against antibiotic-associated diarrhea [48]. Nisin, a peptide produced by *L. lactis*, was investigated for its efficacy against *Clostridium difficile-*associated diarrhea and found to be more effective than Vancomycin and Metronidazole [49]. Besides, *L. raffinolactis* can breakdown α-galactosides, which is a sugar responsible for inducing flatulence and diarrhea [50]. Evidently, our study supports the previous studies, along with indicating that our isolates, especially *Lactococcus lactis* and *Lactococcus piscium*, have the potential to be promising probiotic agents for the prevention of antimicrobial-associated diarrhea.

## Conclusion

In conclusion, the experimental results showed that isolated five selected species could tolerate inhibitory substance bile salt at 1-3 %. The suitable temperature for their growth is showed 37 °C and pH-4.5, but *Lactobacillus delbrueckii* and *Lactobacillus plantarum* grew at temperature 45 °C (data not shown). In comparison to Loperamide, we found that our isolates, especially *Lactococcus lactis* and *Lactococcus raffinolactis*, have the same effects on the rat. Growth inhibition against selected pathogens and resistance against various antibiotics suggests that our isolates may have the potentiality as therapeutics.

## Acknowledgment

We thank the Bangladesh Council of Scientific and Industrial Research (BCSIR) for their logistic support and collaboration.

